# Neural correlates of an illusionary sense of agency caused by virtual reality

**DOI:** 10.1101/2022.09.21.508963

**Authors:** Yiyang Cai, Huichao Yang, Xiaosha Wang, Ziyi Xiong, Simone Kühn, Yanchao Bi, Kunlin Wei

## Abstract

Sense of agency (SoA) is the sensation that self-actions lead to ensuing perceptual consequences. The prospective mechanism emphasizes that SoA arises from motor prediction and its comparison with actual action outcomes, while the reconstructive mechanism stresses that SoA emerges from retrospective causal processing about the action outcomes. Consistent with the prospective mechanism, motor planning regions were identified by neuroimaging studies using the temporal binding effect, a behavioral measure often linked to implicit SoA. Yet, temporal binding also occurs during passive observation of another’s action, lending support to the reconstructive mechanism, but its neural correlates remain unexplored. Here, we employed virtual reality (VR) to modulate such observation-based SoA and examined it with functional magnetic resonance imaging. After manipulating an avatar hand in VR, participants passively observed an avatar’s “action” and showed a significant increase in temporal binding. The binding effect was associated with the right angular gyrus and inferior parietal lobule, which are critical nodes for inferential and agency processing. These results suggest that the experience of controlling an avatar may potentiate inferential processing within the right inferior parietal cortex and give rise to the illusionary sense of agency without voluntary action.

## Introduction

Sense of agency (SoA) is the sensation that self-initiated actions influence the external environment. We implicitly experience the feeling of the connection between our action and the resulting consequence and attend to its disruption only when the actual action feedback conflicts with our expected consequences. As an integral part of self-consciousness, SoA enables one to feel fluent control over one’s surroundings (Haggard 2017), distinct from others (Kahl and Kopp 2018), and responsible for one’s own actions (Haggard and Tsakiris 2009). A clear understanding of the computations underlying SoA is still lacking but two major mechanisms are currently attested. The prospective mechanism emphasizes that SoA is based on a predictive process in the motor system and a comparative process for comparing the predicted and actual action feedback (Frith et al. 2000; Gallagher 2000; Wolpert and Ghahramani 2000; Haggard 2005). The predictive process uses the efference copy of the current motor commands to generate expectations of action consequences. A mismatch between the prediction and actual sensory feedback can disrupt the otherwise fluent SoA. On the other hand, the reconstructive mechanism emphasizes that SoA arises from retrospective explanations of sensory feedback after movement (Wegner and Wheatley 1999; Wegner 2003; Buehner and Humphreys 2009). This inferential sensemaking process evaluates the action feedback and its contingency with prior intentions and goals, reconstructing the causal links between them. While both mechanisms depend on the processing of sensory feedback, they differ in predictive aspects of motor control: the prospective mechanism necessitates the forward model of motor control, i.e., the sensory prediction of action consequence, while the reconstructive mechanism does not rely on the forward model but necessitates post-movement inferential processing of action feedback.

Previous neuroimaging studies typically modulate the magnitude of SoA by either manipulating the authorship of the action, e.g., externally moving people’s effector to generate passive “actions” (Balslev et al. 2006; Tsakiris et al. 2010; Kuhn et al. 2013; Straube et al. 2017; van Kemenade et al. 2017, 2019; Uhlmann et al. 2020; Zapparoli et al. 2020), or perturbing the sensory feedback of the movement or its outcome by implementing temporal and spatial discrepancies (Farrer and Frith 2002; Farrer et al. 2003, 2008; Leube, Knoblich, Erb, and Kircher 2003; Leube, Knoblich, Erb, Grodd, et al. 2003; Matsuzawa et al. 2005; Balslev et al. 2006; David et al. 2007; Schnell et al. 2007; Spengler et al. 2009; Yomogida et al. 2010; Nahab et al. 2011; Chambon et al. 2013; Kuhn et al. 2013; de Bezenac et al. 2016; Sasaki et al. 2018; Kikuchi et al. 2019; van Kemenade et al. 2019; Di Plinio et al. 2020; Ohata et al. 2020; Uhlmann et al. 2020; Zapparoli et al. 2020). Such contrasts between voluntary action and perturbed action revealed neural correlates of SoA in extensive cortical areas such as frontal, parietal, temporal, and insula cortices and subcortical regions such as the cerebellum and striatum (Haggard 2017; Seghezzi et al. 2019; Charalampaki et al. 2022).

Temporal binding (TB), adopted by many as an indicator of implicit SoA (Haggard et al. 2002; Moore and Obhi 2012; Haggard 2017; Tanaka et al. 2019), refers to the fact that people’s timing judgment of an action (e.g., a key press) and its delayed outcome (e.g., a beep sound or flash) are biased towards each other whenever the movement is voluntary as compared to involuntarily made (e.g., the finger pushed by others or triggered by transcranial magnetic stimulation (TMS; Haggard et al. 2002). Studies on the neural substrate underlying TB have highlighted the activity in a brain network including the pre-supplementary motor area (pre-SMA; Kuhn et al. 2013) and dorsal parietal cortex (Seghezzi and Zapparoli 2020; Zapparoli et al. 2020). In fact, modulating the activity over the pre-SMA by repetitive TMS selectively at the motor planning phase affects the binding effect (Zapparoli et al. 2020), with similar findings by the use of transcranial direct current stimulation and theta-burst TMS (Moore et al. 2010; Cavazzana et al. 2015). Given that the pre-SMA is a key structure for preparing and initiating voluntary actions (Cunnington et al. 2003), these findings have been used as neural support for the prospective mechanism of SoA.

Behavioral studies, however, highlighted that motor planning and execution are not necessary for temporal binding, given that it can be elicited without voluntary action (Buehner and Humphreys 2009; Buehner 2012; Poonian and Cunnington 2013; Dewey and Knoblich 2014; Poonian et al. 2015; Borhani et al. 2017; Kong et al. 2017; Vastano et al. 2018; Suzuki et al. 2019). The temporal binding effect can be generated by merely observing another human’s or even a machine’s causal action (i.e., a key press), while observing a non-causal event (i.e., an LED flash) could not (Buehner 2012). These observation-elicited binding effects thus support the reconstructive mechanism, which conceptualizes SoA as a consequence of post hoc inference after movements (Wegner and Wheatley 1999; Wegner 2003; Buehner and Humphreys 2009; Desantis et al. 2011). Given the behavioral evidence, many researchers propose that both predictive and retrospective processes contribute to the manifestation of SoA (Moore and Obhi 2012). However, the neural evidence supporting observation-elicited implicit SoA and thus the reconstructive mechanism is currently lacking.

Here we used virtual reality (VR) to modulate people’s SoA, which is measured by a modified temporal binding task *without* requiring them to execute movements, and examined whether its neural correlates were specifically tied to the inferential processing of action feedback rather than to motor planning and execution. Our recent behavioral study showed that after controlling an avatar in a first-person perspective in VR for a brief period, people increased temporal binding when passively observing an avatar’s “action” (Kong et al. 2017). This “embodiment” effect was thus caused by the prior experience of controlling the avatar since the temporal binding was unchanged for people who experienced the identical VR environment but without controlling the avatar. Hence, such a VR setting would allow us to modulate the implicit temporal binding and reveal its related neural changes when no voluntary action is engaged. We hypothesized that if the change in binding involved motor processes, we should find its neural correlates in sensorimotor regions, especially those planning areas (e.g., pre-SMA and SMA proper) implicated in motor intention and planning (Sperduti et al. 2011; Seghezzi et al. 2019). Alternatively, if post-movement inferential processes contributed heavily to the binding effect, we should observe its neural correlates in the regions outside the frontal motor areas. The targeted areas included posterior parietal areas that had been attributed to causal inference and action awareness (Wende et al. 2013; Renes et al. 2015; Haggard 2017). In particular, inferior parietal regions deserved special attention since direct stimulation of these regions induced subjective experiences of intending to move or even increased (false) reports of movements that were not objectively measured (Desmurget et al. 2009).

Another venue of our study was that our VR manipulation enabled us to examine the neural basis of SoA over a virtual body. VR experience could change people’s self-consciousness (Slater et al. 2009; Banakou and Slater 2014), SoA included (Banakou and Slater 2014; Kokkinara et al. 2016; Padrao et al. 2016; Nierula et al. 2021). However, previous neuroimaging studies focused on the sense of bodily ownership (Bach et al. 2012; Bekrater-Bodmann et al. 2014; Pamplona et al. 2022) and self-localization (Ionta et al. 2011; Lenggenhager et al. 2011). The neural substrate underlying SoA over a virtual body is still understudied (but see Nahab et al. 2011; Padrao et al. 2016; Limanowski et al. 2017, 2018), especially for the temporal binding effect. Furthermore, previous neural studies on SoA over virtual body typically contrast conditions with different levels of spatiotemporal mismatch between virtual and actual actions. Our paradigm, instead, enables us to examine the neural correlates of embodying an avatar by contrasting before and after a VR experience.

## Materials and Methods

### Participants

Our study recruited 48 college students as paid volunteers. Half of the participants were randomly assigned to the experimental group and half to the control group. Both groups were exposed to a VR environment, but only the experimental group viewed an avatar hand in VR. Three participants from the control group were excluded from data analysis, one for excessive head motion (> 2 mm maximum translation or 2° rotation), and two for technical failure (details in Procedures), leaving 24 participants in the experimental group (age: *M* ± *SD* = 23.57 ± 2.39 years, 12 females) and 21 participants in the control group (age: *M* ± *SD* = 22.18 ± 2.76 years, 13 females). Power analysis was conducted based on the reported effect size in our previous study with a similar design (Cohen’s *f* = .5 for the interaction effect in a two-way mixed ANOVA; Kong et al. 2017), and indicated that a sample size of n = 14 per group would lead to a power of 0.9 with an α level of 0.05 (G*Power 3.1; Faul et al. 2007). The two groups were matched on age (*t*_43_ = 1.81, *p* = .078) and gender (*X*^2^_1_ = .643, *p* = .423). All participants were right-handed, had normal or corrected-to-normal vision, and reported no neurological diagnoses. The experiment was conducted according to the principles of the Declaration of Helsinki and was approved by the Ethical Committee of the School of Psychological and Cognitive Sciences at Peking University.

### Procedures

#### Experimental procedure overview

Each participant went through three consecutive phases of the experiment, i.e., a pre-test, VR exposure, and a post-test. The pre-test and post-test were carried out in the MRI scanner (each lasting about 35 minutes), in which participants performed the classical temporal binding task (see “Temporal binding task” below) and a hand laterality judgment task (Ferri et al. 2012) in a sequel; The laterality task was designed to study questions unrelated to the purpose of the current investigation and was not reported here. After the pre-test, the participants walked into the waiting room next to the scanning room to receive VR exposure for about 30 minutes. During VR exposure, participants wore a head-mounted display (HMD, HTC Vive Pro) and a motion-tracking glove (Noitom Hi5 VR Glove) to perform four gamified motor tasks (see “VR exposure” below). The experimental group could view an avatar hand, whose motion spatially and temporally matched with that of their actual right hand; by controlling the avatar hand for these goal-directed movements, VR exposure would enable participants to embody the avatar hand. The control group was never given a chance to see the avatar hand and performed the same motor tasks. After finishing the VR exposure, the participants removed the HMD and the motion-tracking glove, and walked with their eyes closed to the scanning room under the guidance of the experimenter. They were instructed not to open their eyes until they were properly positioned in the scanner to get ready for the post-test. This procedure was employed to minimize the visual experience of the real settings and to preserve the effect of VR exposure. After the post-test, participants re-entered the waiting room and were asked to evaluate their subjective sense of embodiment during the post-test by means of questionnaires.

Participants performed four VR motor tasks: the gesture-imitation task: bending right-hand fingers to match a target gesture, shown by the distant avatar hand; the bubble-poking task: poking the bubble with the right index finger; the cube-picking task: picking target cubes that are specified by color or shape in instructions; the pad-tapping task: memorizing a multi-digit number before it disappears and then recalling it by tapping on a keypad. The experimental group viewed the avatar hand (shown here), but the control group did not. (**C**) Temporal binding effects were quantified by perceptual shifts evoked by the avatar’s movement. The average binding effects in the pre-test and post-test are shown for the two groups separately.

#### Temporal binding task

The temporal binding task was a replicate of the same task in our previous study, though it had been performed with the HMD earlier (Kong et al. 2017). In brief, participants judged the timing of an auditory stimulus with the aid of a Libet clock projected in the MRI scanner in keeping with the temporal binding task previously performed outside the VR (Haggard et al. 2002).

The task involved temporal judgments of tones in two types of trials, baseline and operant trials. For each operant trial (top panel in Fig. 1a), the Libet clock started to rotate clockwise from a random location. After a random interval of 2560 - 5120 ms, the right avatar hand pressed the white button. A tone would be presented 250 ms later (100 ms in duration). Note that the participant was required to refrain from any movement during the stimulus presentation. The clock hand kept rotating after the tone for a random duration of 1280 - 2560 ms, then reset its position to 12 o’clock. Participants were required to report the location of the clock hand when the tone was perceived by pressing the left and right keys using the left middle and index finger to move the clock hand clockwise and counterclockwise, respectively. They then confirmed their estimated position by pressing the key with the right index finger. The report was self-paced. The next trial started upon the confirmation. The baseline trial (bottom panel in Fig. 1a) was identical to the operant trial, except that the tone was presented without the button press, i.e., the avatar’s hand remained static all the time. The mean trial duration was 12.06 s (SD: 2.98 s). Each run contained 20 trials, lasting about 6-8 minutes. Both the pre-test and the post-test were comprised of two operant runs and two baseline runs, whose orders were randomized and counterbalanced between participants.

**Fig. 1.**
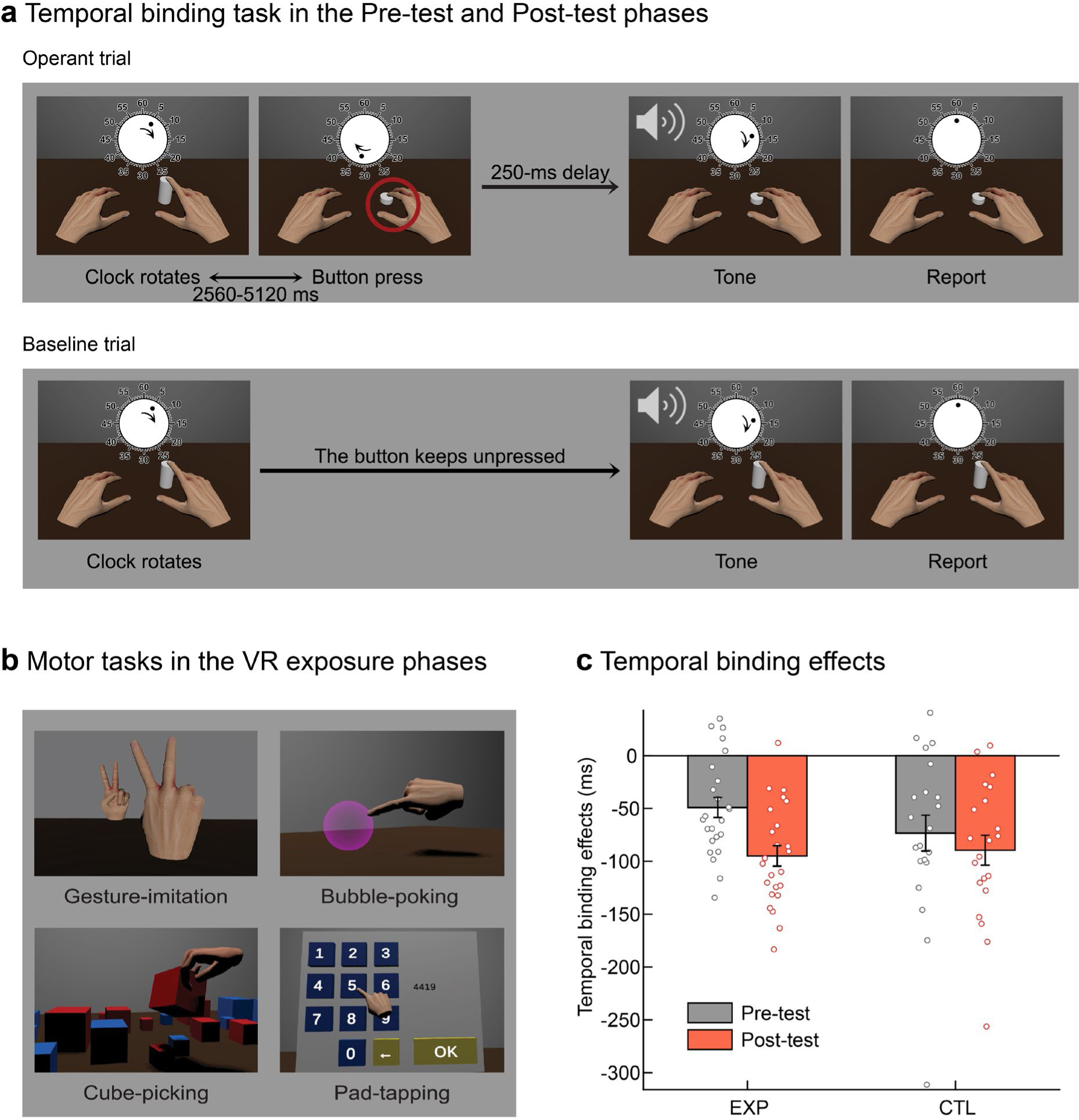
Illustrations of behavioral tasks and the observed temporal binding effects. (**A**) Graphical illustration of an operant trial (top panel) and a baseline trial (bottom panel) in the temporal binding task. In both conditions, participants were required to report the location of the clock hand when the tone was perceived. Throughout the baseline trials, the avatar hands on the screen kept stationary. But in the operant trials, its right index finger “pressed” a white button 250 ms prior to the tone. Participants’ real hands kept unmoved before making temporal judgements. (**B**) Scenes of motor tasks in the VR exposure phase.

Participants familiarized themselves with the temporal binding task before the formal experiment by conducting four operant trials and four baseline trials outside the scanner. Due to technical failure, two control participants underwent one operant run and three baseline runs in one of the two phases, and were excluded from data analysis.

All the visual stimuli were back-projected onto a translucent screen located inside the scanner (resolution: 1024×768; refresh rate: 60Hz; viewing distance about 90 cm; Fig. 1a). The avatar hand was the one that the participants visually controlled during the VR exposure outside the scanner but fitted for rendering in the two-dimension display in the scanner. The clock had a 10-pixel-long hand, which rotated with a period of 2560 ms per circle. The clock face (radius = 110 pixels) was marked with conventional “1 min” intervals. The participants wore a pair of headphones to receive the auditory stimuli and to reduce noise. We adjusted the volume of the auditory stimulus on the individual basis to ensure that they could hear the stimuli clearly and comfortably in the scanner. The rendering was controlled by a customized program coded using Psychtoolbox (3.0.16) implemented in MATLAB (R2016b, MathWorks).

#### VR exposure

During VR exposure that was sandwiched between the pre-test and the post-test, participants wore the HMD (HTC Vive Pro) with a dual-display giving a 110° field of view and a resolution of 1080 × 1200 per eye. In the VR scene, they could see a virtual desk located at approximately the same location as the physical desk at which they sat. The motion tracking glove sampled their right-hand motion at a sampling rate of 100 Hz to control a right avatar hand in the VR environment. The motion of the avatar hand was always temporally and spatially congruent with the real right hand. For participants from the experimental group, the avatar hand resembled their own right hand in size and skin tone, which were adjusted before the experiment. For control participants, no avatar hand was shown, and all the VR tasks were performed with the “invisible” hand. The games were designed to provide a sensorimotor experience of controlling the avatar’s hand to interact with the virtual environment (Fig. 1b), including the gesture-imitation task, the bubble-poking task, the cube-picking task, and the pad-tapping task (customized by using Unity3D 2019.3.14f1).

In the gesture-imitation task, a left-hand avatar was displayed in front of the participants at their eye height. This avatar hand would present a hand gesture randomly selected from 20 possible gestures, e.g., a closed fist, a thumb-up, and a V-sign. The participant was required to replicate the same gesture with their right avatar hand. Once the gesture was reproduced, the next target gesture would appear until all 40 trials (two repetitions for each different gesture) were successfully finished. In the bubble-poking task, a transparent bubble would appear at a random location on the desk, moving at a random speed and direction. Participants were asked to poke the bubble with the index finger of the right avatar hand before it fell off the desk. After a bubble was poked or dropped off the desk, the next bubble would appear. This task lasted until 60 target bubbles were shown and poked. In the cube-picking task, forty cubes with different sizes, colors (red or blue), and labeled numbers (1 or 2) were placed randomly on the virtual desk. Participants were required to control the avatar hand to grab specific cubes and drop them on the floor. Participants performed this task three times with different verbal instructions each time, including “Pick all the red cubes and drop them on the floor”, “Pick all the cubes labeled as number 1 and drop them on the floor”, and “Pick all the red cubes labeled as number 1 and drop them on the floor”. In the pad-tapping task, a multi-digit number would appear for 2-4 s. Participants were required to memorize the number and then replicate the number by tapping on a virtual number pad. Forty numbers, ranging from 3-digit to 10-digit, were presented in random order for each participant. Each of the four VR tasks lasted for about 6-7 minutes, resulting in a total VR exposure duration of about half an hour.

#### Questionnaire

We assessed the subjective feeling of embodiment using questionnaires after the post-test. The questions were identical to the ones used in our previous study on VR embodiment (Kong et al. 2017). The items were designed to assess explicit SoA and sense of ownership (SoO) or to control for possible response biases by using the reversed control items (Table S1).

### Behavioral analysis

To examine whether the VR exposure could enhance participants’ implicit SoA associated with an avatar hand, we compared the temporal binding effect using a two-way mixed-design analysis of variance (ANOVA) with Phase as the within-group factor (pre-versus post-test) and Group as the between-group factor (experimental versus control group). Each trial yielded a perceptual error of temporal judgment, quantified as the difference between the reported and the actual onset of the tone. The temporal binding effect was operationally defined as the difference in perceptual error, i.e., a perceptual shift, between the operant condition and the baseline condition. A negative perceptual shift indicated the temporal binding effect.

The present study focused on the implicit SoA, measured using the behavioral temporal binding effects mentioned above, not the explicit measures of embodiment from questionnaires, as our previous behavioral study showed that explicit SoA did not change after brief VR exposure (Kong et al. 2017). As shown in Supplementary Materials, our results also revealed no significant group differences in the explicit measures of embodiment from questionnaires. We also failed to observe a significant correlation between temporal binding or binding changes and these explicit measures. Results and possible implications of explicit ratings are detailed in the Supplementary Materials.

### MRI acquisition

All functional and structural MRI data were acquired on a Siemens 3T Prisma scanner with a 64-channel head-neck coil at the Center for MRI Research, Peking University. High-resolution functional images were acquired using a multi-band echo-planar sequence (62 axial slices, repetition time (TR) = 2000 ms, echo time (TE) = 30 ms, flip angle (FA) = 90°, field of view (FOV) = 224 × 224 mm^2^, matrix size = 112 × 112, slice thickness = 2.0 mm, voxel size = 2 × 2 × 2 mm^3^, multi-band factor = 2). High-resolution 3D T1-weighted anatomical images were acquired before the pre-test using the magnetization-prepared rapid-acquisition gradient-echo sequence (192 sagittal slices, TR = 2530 ms, TE = 2.98 ms, inversion time = 1100 ms, FA = 7°, FOV = 224 × 256 mm^2^, matrix size = 224 × 256, interpolated to 448 × 512, slice thickness = 1.0 mm, voxel size = 0.5 × 0.5 × 1 mm^3^).

### MRI analysis

fMRI data were preprocessed using Statistical Parametric Mapping software (SPM12; http://www.fil.ion.ucl.ac.uk/spm12/) and analyzed using the toolbox for Data Processing & Analysis for Brain Imaging (DPABI, Version 6.2; http://rfmri.org/DPABI; Yan et al. 2016) implemented in MATLAB (R2021a, MathWorks). After preprocessing, the statistical analyses were conducted in R (Version 4.3.1). Bayesian analysis was conducted in JASP (Version 0.18.1; https://jasp-stats.org/). All of the surface-view brain results and ROIs were visualized with the BrainNet Viewer (Version 1.7; http://www.nitrc.org/projects/bnv/; Xia et al. 2013).

#### Data preprocessing

Functional images were preprocessed using SPM12. For each participant, the first five volumes of each functional run were discarded. The remaining images were corrected for slice timing and head motion and spatially normalized to Montreal Neurological Institute (MNI) space via unified segmentation (resampling into 2 × 2 × 2 mm^3^ voxel size). One participant from the control group showed excessive head motion (> 2 mm or 2°) and was excluded from data analysis. The resulting images were spatially smoothed using a 6-mm full-width half-maximum (FWHM) Gaussian kernel for univariate contrast analyses.

#### Generalized linear model

At the first (individual) level, preprocessed functional images of the pre-test and post-test temporal binding tasks were modeled in a generalized linear model (GLM). As the conditions (operant or baseline) were presented in separate runs (with the condition order randomized and counterbalanced across participants), for each trial we regarded the trial onset as a tighter baseline for temporal judgment at tone onset and thus included two regressors corresponding to the two onsets for each run. Each regressor was convolved with the canonical hemodynamic response function (HRF). The GLM also included six predictors of head motion parameters for each run. The high-pass filter was set at 128 s. After model estimation, whole-brain contrast images of operant condition (tone versus start) versus baseline (tone versus start) in the pre-test and post-test were calculated for each participant for further analyses.

#### Regions of interest (ROI) definition

Two types of ROI were defined to examine their possible involvement in VR-induced changes of temporal binding effects. *(1) Literature-based ROIs* were defined on the basis of a previously published meta-analysis on SoA (Seghezzi et al. 2019). We included all the seven regions (Table 1 and Fig. 2a) reported in the meta-analysis, including clusters associated with self-SoA (SoA attributed to self; their Table 1) and with external-SoA (SoA attributed to others; their Table 3), for completeness, as both were related to SoA processing. The ROIs were defined as 8-mm-radius spheres centering on each MNI coordinate. *(2) Task-based ROIs* were functionally defined using our pre-test data. As shown in our behavioral results, in the pre-test, temporal bindings could be elicited merely by observation of “avatar hand” actions. We thus defined the observation-elicited SoA ROIs by contrasting operant versus baseline conditions across all participants, which followed the behavioral operationalization of the temporal binding effects. Only one cluster was found at the threshold of voxel-level one-tailed *p* < .001 and cluster-level family-wise error (FWE) corrected *p* < .05 (Table 1 and Fig. 2b).

**Fig. 2.**
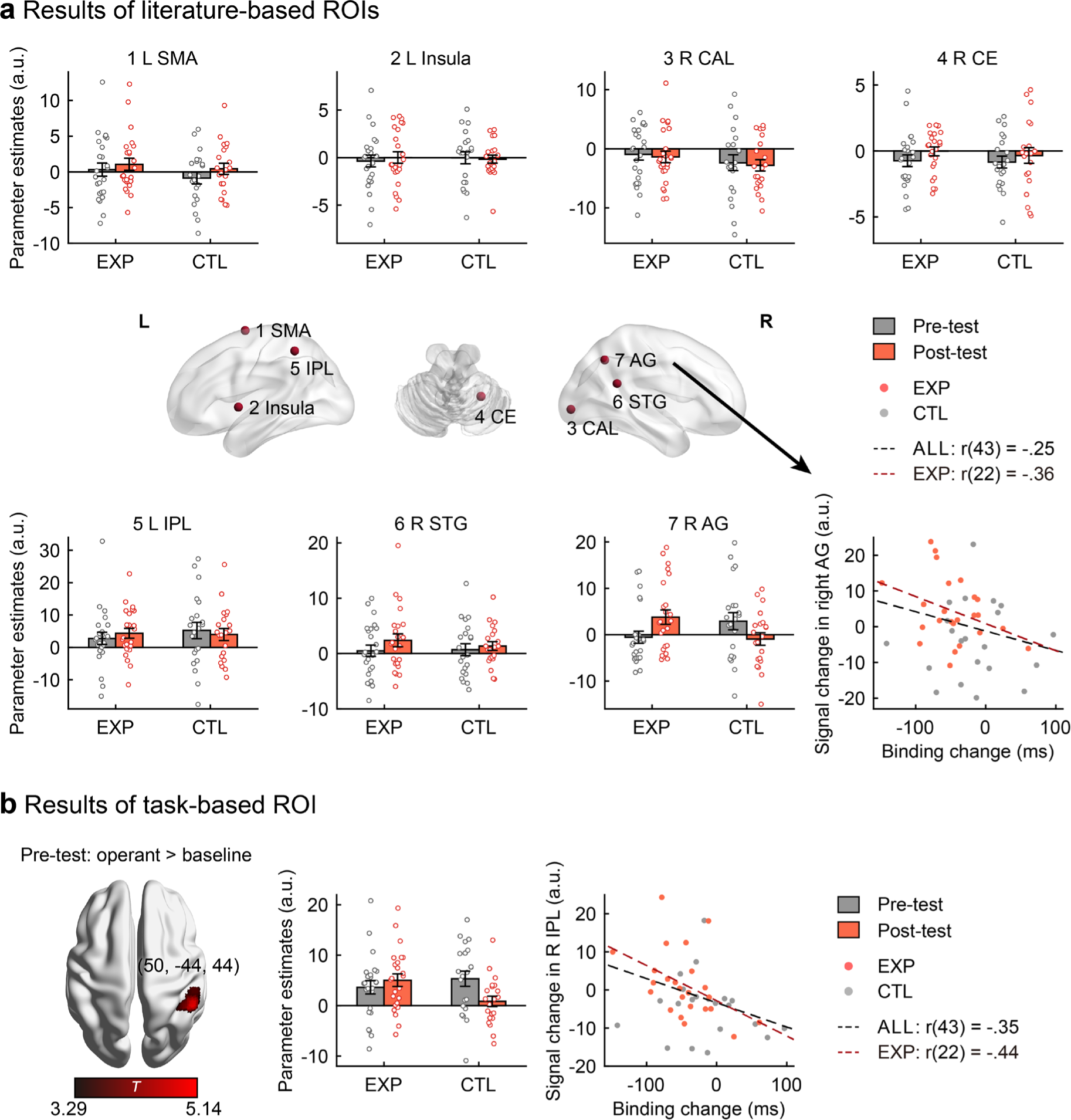
ROI-level results. (**a**) Literature-based ROI results. Seven ROIs were defined on the basis of a previously published meta-analysis (Seghezzi et al. 2019). The locations of the ROIs were displayed in the center penal (details in Table 1). The bar plots showed the mean beta values of operant versus baseline in different phases (gray bars for pre-test and red bars for post-test) and different groups (EXP = experimental group, CTL = control group), with the scatter points showing the individual data. The right AG yielded a significant Phase× Group interaction effect. The lower right scatter graph depicted the association between the temporal binding changes and the VR exposure-related signal changes in the right AG. The regression lines for all participants (black dashed line) and the experimental group alone (red dashed line) were displayed. (**b**) Results of the task-based ROI, defined as regions showing stronger activation in the operant condition relative to the baseline in the pre-test for all participants. Left panel: Only one cluster centered at the right IPL at the threshold of voxel-level one tailed *p* < .001 and cluster-level FWE corrected *p* < .05 (details in Table 1). Center penal: The bar plot showed the mean beta values of operant versus baseline in different phases (gray bars for pre-test and red bars for post-test) and different groups (EXP = experimental group, CTL = control group), with the scatter points showing the individual data. Right penal: the scatter graph depicted the association between the temporal binding changes and the VR exposure-related signal changes in the right IPL cluster. The regression lines for all participants (black dashed line) and the experimental group alone (red dashed line) were displayed.

**Table 1.** Summary of the ROI-level results.

| Location |  | Coordinates |  |  | Phase × Group |  |  |  | Correlation with binding changes |  |  |  |
| --- | --- | --- | --- | --- | --- | --- | --- | --- | --- | --- | --- | --- |
|  |  | (MNI, mm) |  |  | interaction effects |  |  |  | ALL |  | EXP |  |
| Area | H | x | y | z | $F_{1,43}$ | p | Cohen's $f$ | $BF_{incl}$ | $r_{43}$ | p | $r_{22}$ | p |
| **Literature-based ROIs** |  |  |  |  |  |  |  |  |  |  |  |  |
| SMA | L | -7 | -4 | 69 | .11 | .745 | .050 | .302<sup>H0</sup> | .05 | .737 | -.18 | .401 |
| Insula | L | -41 | 2 | 1 | .13 | .716 | .056 | .314<sup>H0</sup> | -.09 | .566 | -.36 | .083 |
| CAL | R | 18 | -90 | -1 | < .001 | .993 | .001 | .325<sup>H0</sup> | .08 | .607 | .07 | .724 |
| CE | R | 24 | -53 | -27 | .08 | .783 | .042 | .294<sup>H0</sup> | .03 | .827 | -.11 | .601 |
| IPL | L | -46 | -48 | 51 | .58 | .450 | .116 | .407 | -.20 | .195 | -.37 | .077 |
| STG | R | 54 | -49 | 22 | .39 | .538 | .095 | .349 | -.06 | .697 | -.29 | .168 |
| AG | R | 45 | -60 | 43 | 7.67 | .008** | .422 | 13.546 | -.25 | .092 | -.36 | .087 |
| **Task-based ROI** |  |  |  |  |  |  |  |  |  |  |  |  |
| IPL | R | 50 | -44 | 44 | 5.29 | .026* | .351 | 3.708 | -.35 | .017* | -.44 | .029* |

Notes: H = Hemisphere, L = left, R = right; SMA = Supplementary motor area, Insula = Posterior insula, CAL = Calcarine scissure, IPL = Inferior parietal lobule, AG = Angular gyrus, STG = Superior temporal gyrus, CE = Cerebellum; <sup>H0</sup>: moderate evidence in support of the null hypothesis of no interaction effect ( $BF_{incl} < .33$ ); $\because .05 < p < .10$ , \*: $.01 < p < .05$ , \*\*: $p < .01$ .

**Table 2.** Summary of the whole-brain contrasts.

| Location |  | Coordinates (MNI, mm) |  |  | k<sub>E</sub> | Peak *t* | Cluster *p*<sub>FWE-corr</sub> |
| --- | --- | --- | --- | --- | --- | --- | --- |
| Area | H | x | y | z |  |  |  |
| Experimental group (n = 24): binding effects (operant vs. baseline) in the post-test |  |  |  |  |  |  |  |
| (Threshold: voxel-level one-tailed *p* < .001 and cluster-level FWE corrected *p* < .05) |  |  |  |  |  |  |  |
| IFG | R | 52 | 18 | 28 | 386 | 7.15 | <.001 |
| IPL | L | -40 | -38 | 52 | 297 | 6.64 | <.001 |
| Precuneus | L | -12 | -70 | 54 | 83 | 6.19 | .036 |
| Precuneus | R | 8 | -66 | 52 | 350 | 5.59 | <.001 |
| IPL | R | 46 | -40 | 34 | 256 | 5.57 | <.001 |
| SFG | L | -28 | -4 | 60 | 89 | 4.78 | .026 |
| Control group (n = 21): binding effects (operant vs. baseline) in the post-test |  |  |  |  |  |  |  |
| (Threshold: voxel-level one-tailed *p* < .001 and k > 20) |  |  |  |  |  |  |  |
| IPL/SMG | L | -52 | -38 | 34 | 23 | 5.32 | .849 |
| MTG | R | 62 | -54 | 8 | 23 | 4.83 | .849 |
| Between-group contrast (Exp vs. Ctl) of the VR exposure-related effects |  |  |  |  |  |  |  |
| (Threshold: voxel-level one-tailed *p* < .001 and k > 20) |  |  |  |  |  |  |  |
| IFG | R | 54 | 16 | 28 | 55 | 4.45 | .262 |
| AG | R | 38 | -66 | 54 | 20 | 3.70 | .941 |

Notes: H = Hemisphere, L = left, R = right, $k_E$ = cluster size (vx).

#### ROI-level Phase × Group interaction effects

To examine the neural basis of VR-induced changes of implicit SoA, we examined the Phase × Group interaction effects at the ROI level. In each ROI we defined above, the averaged beta values of the operant condition versus baseline in different phases (pre-or post-test) were extracted for each participant. These values were examined using two-way mixed ANOVAs, with Phase as the within-group factor (pre-or post-test) and Group as the between-group factor (experimental or control group). For ROIs revealing insignificant interaction effects (*p* > .05), Bayesian ANOVAs were conducted by JASP (0.18.1) with the default setting. The effects of the components were calculated by comparing across the matched models. The model-averaged inclusion Bayesian factor (*BF*_incl_) of the Phase × Group interaction component was used to quantify the evidence in favor of the null hypothesis (no interaction effect), with .33 < *BF*_incl_ < 1 as anecdotal evidence, .1 < *BF*_incl_ < .33 as moderate evidence, and *BF_inclu_* < .1 as strong evidence (van den Bergh et al. 2020).

#### Whole-brain univariate analyses

In the task-based ROI analysis, a one-sample *t-*test of the contrast images obtained in the pre-test for all participants was carried out to localize the regions underlying the temporal binding effects. Similar *t*-tests for the post-test were conducted for the experimental and control groups, respectively, to localize similar regions after VR exposure. Besides, the VR exposure-related neural activities were quantified by subtracting the whole-brain contrast images of the pre-test (operant versus baseline) from the post-test (operant versus baseline) for each participant. A two-sample *t-*test was then conducted to compare the two groups, which is equivalent to the Phase × Group interaction effects. All whole-brain results were thresholded at voxel-wise one-tailed *p* < .001, cluster-level FWE-corrected *p* < .05. For the contrasts that did not survive the cluster-level FWE correction, clusters exceeded 20 voxels (voxel-wise threshold at one-tailed *p* < .001) were reported.

#### Correlation analyses between fMRI and behavioral measures

To further examine whether VR-induced neural changes were associated with VR-induced behavioral changes, we computed the Pearson correlation between behavioral changes in temporal binding effects and the neural signal changes in the ROIs showing significant Phase × Group interaction effects and at the whole-brain level. To rule out the possibility that the correlations may simply arise from group differences, we also carried out the correlation in the experimental group alone.

## Results

### Behavioral results: the temporal binding effect

Before the VR exposure (during the pre-test phase), both groups of participants showed a moderate level of temporal binding effect (Fig. 1C). In the baseline condition, when the avatar hand remained stationary, the temporal judgment error amounted to an average of −26.23 ms (*SE* = 13.86 ms, same below) and - 45.89 (17.75) ms for the experimental and control groups, respectively. In the operant condition with the avatar hand suddenly pressing the button before the tone, the judgment error was −75.27 (16.10) ms and - 119.18 (24.99) ms for the experimental and control groups, respectively. Thus, in the pre-test, both groups showed a negative perceptual shift in timing judgment of the tone, an indicator of temporal binding. Notably, between-group comparison of the temporal binding effects in the pre-test was not significant (*t*_43_ = 1.29, *p* = .206, Cohen’s *d* = .38), suggesting that the magnitudes of the binding effects in the pre-test were matched across groups.

Crucially, after the VR exposure in the post-test, the experimental group increased their binding effects by 45.80 ms (8.69) with the baseline and operant error of −5.16 (14.08) ms and −100.01(18.23) ms, respectively. The control group’s binding effects remained largely unchanged (differed from the pre-test by −16.17 ± 11.68 ms, with a baseline error of −26.73 ± 14.79 ms and an operant error of −116.20 ± 23.32 ms). This group difference was confirmed by a mixed ANOVA (within-group factor: Phase; between-group factor: Group): both the interaction effect (*F*_1, 43_ = 4.27, *p* = .045, Cohen’s *f* = .32) and the main effect of Phase (*F*_1, 43_ = 18.68, *p* < .001, Cohen’s *f* = .66) reached significance, while the main effect of Group was not significant (*F*_1, 43_ = .33, *p* = .567, Cohen’s *f* = .09). Post-hoc comparisons indicated that the experimental group showed enhanced binding effects in the post-test (−94.85 ± 11.46 ms) compared to the pre-test (−49.04 ± 12.89 ms; *t* = 4.68, Bonferroni-corrected *p* < .001, Cohen’s *d* = .97), while the same effect was absent in the control group (*t* = 1.54, Bonferroni-corrected *p* = .130, Cohen’s *d* = .34). These behavioral results and their effect sizes were similar to those in our previous study (Kong et al. 2017), indicating that seeing and controlling an avatar hand enhanced the temporal binding effect.

### Literature-based ROI results

To investigate the possible involvement of the traditional SoA regions (typically induced by self-action, Seghezzi et al. 2019) in our VR-induced SoA effects, we explored whether these regions could show similar Phase× Group interaction effects as the behavioral results above.

Among the seven literature-based ROIs (Table 1 and Fig. 2a), only one ROI centered at the right angular gyrus (AG) yielded a significant interaction effect (*F*_1, 43_ = 7.67, *p* = .008, Bonferroni-corrected *p* = .058, Cohen’s *f* = .42). Post-hoc comparisons indicated that the interaction effect in the right AG was driven by the significantly increased signal after VR exposure with a visible avatar for the experimental group (*t* = 2.14, Bonferroni-corrected *p* = .038, Cohen’s *d* = .44), as well as the marginally significant signal decrease for the control group (*t* = −1.79, Bonferroni-corrected *p* = .081, Cohen’s *d* = −.39). Note that the pre-test signals were comparable across two groups (*t*_43_ = −1.56, *p* = .126, Cohen’s *d* = −.47). For the rest six ROIs showing insignificant interaction effects (all *p*s > .05), Bayesian ANOVAs were conducted to quantify the evidence in favor of the null hypothesis (*H*_0_: no interaction effect). *BF*_incl_ yielded moderate evidence for *H*_0_ in the left pre-supplementary motor area (SMA), the left Insula, the right calcarine scissure (CAL) and the right cerebellum (CE; all *BF*_incl_s < .33), and anecdotal evidence for *H*_0_ in the left inferior parietal lobule (IPL; *BF*_incl_ = .41) and the right superior temporal gyrus (STG; *BF*_incl_ = .35). Thus, although most of regions from the traditional SoA network were not involved in our paradigm, the right AG’s neural response to the implicit SoA was modulated by VR exposure. Importantly, correlation analyses showed that activation changes in this region negatively correlated with behavioral changes in temporal binding at a marginal significance (*r*_43_ = −.25, *p* = .092; lower right panel in Fig. 2a), and this correlation remained for the experimental group alone (*r*_22_ = −.36, *p* = .087). That is, the bigger the activation changes in the right AG (more positive values), the larger the increases in the temporal binding effects (more negative values).

### Task-based ROI results

Unlike the previous imaging studies using voluntary actions to study SoA, we elicited the temporal binding effects merely by observation of “embodied” actions. Thus, we defined another type of ROI to represent the neural responses of our special temporal binding task. As the temporal binding effects were behaviorally calculated as operant errors - baseline errors, the task-based ROI was defined using the contrast “operant versus baseline” in the pre-test for all participants. Only one cluster centered at the right IPL showed significant activation (Table 1 and Fig 2b; peak MNI *xyz*: 50, −44, 44; cluster size = 296 voxels, peak *t*_44_ = 5.14, cluster-level *p*_FWE-corr_ < .001) at the threshold of voxel-level one-tailed *p* < .001 and cluster-level FWE corrected *p* < .05.

The right IPL cluster is anatomically close to the right AG cluster defined in the literature-based ROIs above (16.8 mm between the peak coordinates), and also yielded a similar Phase× Group interaction effect (*F*_1, 43_ = 5.29, *p* = .026, Cohen’s *f* = .35). Post-hoc comparisons indicated that the interaction effect in the right IPL cluster was driven by the significantly decreased signal after VR exposure for the control group (*t* = −4.46, Bonferroni-corrected *p* = .021, Cohen’s *d* = −.53) and no significant signal change for the experimental group (*t* = .80, Bonferroni-corrected *p* = .426, Cohen’s *d* = .17). Note that the pre-test signals were comparable across two groups (*t*_43_ = −.85, *p* = .401, Cohen’s *d* = −.25). Similar to the AG, the right IPL demonstrated a negative correlation between VR-induced neural and behavioral changes. This was observed when data were evaluated for all participants (*r*_43_ = −.35, *p* = .017; lower right panel in Fig. 2b) as well as for the experimental group alone (*r*_22_ = −.45, *p* = .029).

### Whole-brain results

As mentioned above, the binding effects in the pre-test across all participants were found in the right IPL. After VR exposure (visible avatar for the experimental group and invisible for the control group), the binding effects in the post-test were examined for the experimental and control group, respectively. For the experimental group who behaviorally showed enhanced binding effects in the post-test, we observed significant activation in five clusters (Table 1 and top panel of Fig. 3a; voxel-level threshold at one-tailed *p* < .001 and cluster-level FWE correction at *p* < .05), including the right inferior frontal gyrus (IFG), bilateral IPL, bilateral precuneus and left superior frontal gyrus (SFG). Note that the right IPL cluster largely overlaps with the task-based ROI, IPL, defined in the pre-test (Table 1 and Fig. 2b). For the control group, no significant activation was found at the conventional threshold and we observed small clusters in the left inferior parietal lobule and the right middle temporal gyrus at a lenient threshold of voxel-wise *p* < .001, cluster size > 20 voxels (Table 1).

**Fig. 3.**
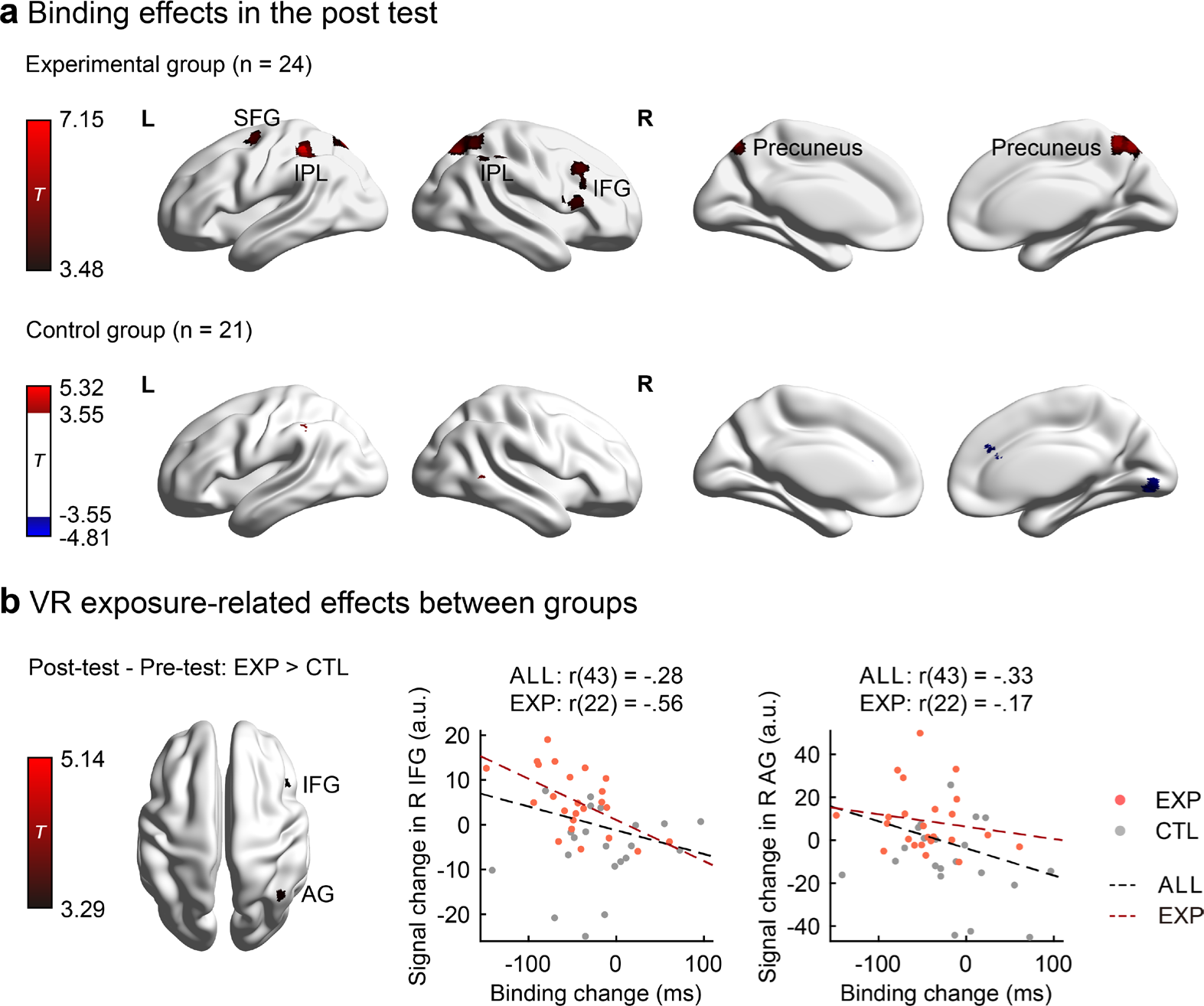
Whole-brain results. (**a**) Whole-brain results of the binding effects in the post-test. Top panel: the experimental group showed activation in the post-test in five clusters (voxel-level threshold at one-tailed *p* < .001 and cluster-level FWE correction at *p* < .05). Bottom panel: no significant activation in the post-test was found for the control group at the same threshold. Clusters exceeding 20 voxels at a voxel-level threshold of one-tailed *p* < .001 were displayed. (**b**) Whole-brain results of the VR exposure-related effects between two groups. Left panel: No significant activation was detected at a voxel-level threshold of one-tailed *p* < .001 and cluster-level FWE correction at *p* < .05. Clusters of the right IFG and right AG exceeded 20 voxels at a voxel-level threshold of one-tailed *p* < .001 and were displayed. Center and Right penal: The scatter graphs depicted the association between the temporal binding changes and the VR exposure-related signal changes in the right IFG and right AG, respectively. The regression lines for all participants (black dashed line) and the experimental group alone (red dashed line) were displayed (EXP = experimental group, CTL = control group).

We then examined the whole-brain Phase × Group interaction effect and found that no regions survived the conventional threshold. At a lenient threshold of voxel-level one-tailed *p* < .001 and cluster size > 20 voxels (Table 1), interaction effects were observed in the right IFG and the right AG (Fig. 3b). The two clusters demonstrated similar interaction patterns in that the experimental group exhibited increased activation (*t*_23_ > 2.07, *p* < .05) and the control group showed reduced activation (*t*_20_ > 2.09, *p* < .05) in the comparison between pre- and post-test data. In terms of behavioral associations, the right AG cluster showed a significantly negative correlation between the binding changes and the signal changes for all participants (*r*_43_ = −.33, *p* = .025), which did not approach significance for the experimental group (*r*_43_ = −.17, *p* = .437); the similarly negative correlation in right IFG was marginally significant for all participants (*r*_43_ = −.28, *p* = .061) and reached significance for the experimental group alone (*r*_43_ = −.56, *p* = .005). Whole-brain correlation analyses did not reveal other significant regions even at a lenient threshold.

## Discussion

We examined the brain activation pattern supporting the changes in sense of agency that were elicited by passive observation of “embodied” avatar actions using the temporal binding paradigm before and after VR exposure. Behaviorally, we replicated our previous findings that controlling an avatar for a short period of time leads to increased binding upon seeing the avatar’s hand action while the actual hand stays immobile (Kong et al. 2017). Extending the previous finding, here we show that this VR embodiment effect persists after exiting the VR setting and when the same avatar hand is then depicted in a two-dimensional display in the fMRI scanner. Our neuroimaging results identified that temporal binding elicited by observation is specifically associated with a cluster centered at the inferior parietal lobule. Accordingly, our ROI analysis with the previously identified SoA network found that the VR enhancement of temporal binding is specifically implicated in the right angular gyrus, not in any motor planning regions such as pre-SMA. The higher the activation changes in the region, the larger the increase in temporal binding (Fig. 2a). As expected, the observed neural correlates for VR-induced SoA are distinct from neural correlates for other VR-induced self-consciousness changes, such as bodily ownership and self-localization. Our findings not only revealed the neural substrate underlying temporal binding elicited by observation, but also provided supporting evidence for the reconstructive mechanism of SoA by showing that temporal binding is subserved by neural regions tied to inferential sensemaking of sensory events.

Previous investigations have extensively examined the activation foci for SoA by contrasting condition pairs with presumably different levels of SoA, i.e., voluntary action versus rest, voluntary action versus stimulus-driven action, actions with higher visuomotor congruency versus those with lower congruency (see a recent review, Seghezzi et al. 2019). Of note, all these contrasting analyses involve voluntary actions. Thus the identified extensive brain network for SoA includes sensorimotor areas, especially those related to intentionality and motor planning (Seghezzi et al. 2019; Zapparoli et al. 2020). For example, the dorsolateral prefrontal cortex is associated with action selection (Khalighinejad et al. 2016), pre-SMA with intentionality (Yomogida et al. 2010; Zapparoli et al. 2018), and temporal binding in particular (Kuhn et al. 2013), and SMA proper with motor planning and initiation (Passingham and Lau 2019). Consistent with the rationale of our task design, all these prefrontal and frontal areas, related to the generation of action before action feedback, returned a null effect in our data (Fig. 1a and Table 1), except the right AG identified by the meta-analysis based on voluntary action-related SoA (Seghezzi et al. 2019).

The temporal binding change elicited by observing an embodied action is not only specifically associated with the right AG, but also associated with a cluster extended to the right IPL. Both are not directly tied to motor planning and movement initiation. This finding supports the reconstructive mechanism of SoA, which emphasizes that the temporal binding is grounded by retrospective causality since the right AG and IPL participate in both SoA and causal processing in general. The right IPL has been one of the most frequently revealed neural correlates of SoA (Farrer et al. 2003, 2008; Schnell et al. 2007; Nahab et al. 2011; Chambon et al. 2013, 2015). Even anosognosia patients who often assert that they performed an action with their paralyzed, immobile limb typically have lesions in the right parietal lobule (Fotopoulou et al. 2008). More importantly, the right IPL is broadly involved in causal processing since the explicit judgment of both physical and social causality relates to neural activations in the right IPL, along with other areas (Wende et al. 2013; Renes et al. 2015). Even seeing a causal event, such as an object collision, elicits more activations in the right IPL than seeing a non-causal event, such as an object launching (Fugelsang et al. 2005).

The AG, similarly implicated by numerous SoA studies, engages in diverse cognitive tasks that require inferential sensemaking. For SoA, meta-analyses have shown that the TPJ, with the right AG included, is related to attributing SoA to others (external SoA, Sperduti et al. 2011) and to the reduction of self-agency (negative SoA, Zito et al. 2020). A recent review also finds AG as a common node for encoding motor intention and SoA (Seghezzi et al. 2019). Beyond agency tasks, the AG has been reliably shown to engage in a wide range of tasks, including reasoning, semantic processing, word reading and comprehension, memory retrieval, attention and spatial cognition, default mode network, and social cognition. A well-received unified theory about the AG’s function, based on the commonality of these tasks, highlights its role in sensemaking, i.e., giving meaning to external sensory information or internal thoughts (Seghier 2013). For instance, the AG engages in the comprehension of speech and written languages (Xu et al. 2005; Obleser and Kotz 2010), especially in solving referential ambiguity (Nieuwland et al. 2007). It also engages in inferring human intention in the theory of mind tasks (Mason and Just 2011). Given its rich anatomical connectivity to widely distributed brain regions, the AG appears suitable for combining diverse information, linguistic and nonverbal (e.g., body movements), prior knowledge (experiences, context, and purpose), and new sensory information, to converge toward plausible accounts of the events. This sensemaking process can be implemented as an active optimization process that combines bottom-up information (i.e., sensory information) with top-down predictions (i.e., prior knowledge and purpose) to minimize surprise according to the free energy principle (Friston 2010). Pertinent to our findings here, the AG is a central region for the inferential sensemaking process in various tasks, among which the agency-related task is an important genre since SoA sets the boundary between self and the external environment (Seghier 2013, 2022).

The involvement of AG and IPL in our observation-based temporal binding is in line with the reconstructive mechanism of SoA (Wegner and Wheatley 1999; Wegner 2003; Buehner and Humphreys 2009; Desantis et al. 2011; Tramacere 2021). Our findings, of course, should not be taken as evidence against the importance of the prospective processing for SoA during voluntary action. There exists extensive behavioral and neural evidence that both the prospective motoric process in motor planning and the retrospective process in outcome evaluation contribute to SoA, though their relative importance depends on available cues and task goals (Moore and Obhi 2012; Synofzik et al. 2013). Even AG, the region we identified as crucial for retrospective processing of SoA, has been shown to monitor signals related to action selection in the dorsolateral prefrontal cortex when participants are required to explicitly report their SoA (Chambon et al. 2013, 2015). Our findings highlight that the brain can indeed invoke SoA-related processing retrospectively when no action is involved.

Our findings can also be viewed as a challenge to the validity of treating temporal binding as an implicit measurement of SoA (Buehner 2012; Kong et al. 2017; Suzuki et al. 2019). Temporal binding with voluntary actions indeed changes according to SoA manipulations, including the aforementioned experimental comparisons between active and passive movements and between congruent and incongruent action feedback. However, temporal binding can also be elicited without action and supported by distinct neural substrates, as shown here. Thus, a parsimonious account of temporal binding posits that it results from top-down causal belief about the timing of sequential events, with or without voluntary action (Hoerl et al. 2020). The belief is about the causal relationship between a movement-related event, not necessarily an intentional action, and a subsequent outcome event. The causal belief is subject to influence from priming, instruction, statistical contingency, and prior belief, which all have been shown to affect temporal binding (Wegner 2003; Aarts et al. 2005; Moore et al. 2009; Ebert and Wegner 2010; Desantis et al. 2011). The causal account thus views temporal binding as a general phenomenon in timing perception and casual belief, beyond a reflection of implicit SoA that has been argued to embed in the motor system. This view resembles the reconstructive mechanism of SoA with its emphasis on inferential processing for sensory events. In this light, the control experience during our VR exposure facilitates the formation of causal belief between avatar “action” and subsequent sensory event, possibly following the principles of ideomotor learning (Elsner and Hommel 2001). The AG and IPL underlie the VR binding effects, and thus might play a role in representing the causal belief. Though risking the curse of reverse inference from neural findings to cognitive processes, our findings support the causal account by showing that the neural substrate underlying our observed VR binding effect involves AG and IPL, important areas supporting the causal inference of sensory events.

Though a quantitative model of SoA is currently lacking, various aspects of temporal binding have been accounted for by probabilistic inference models based on Bayesian cue combination (Moore and Fletcher 2012; Wolpe et al. 2013; Legaspi and Toyoizumi 2019; Lush et al. 2019). The temporal shift of the action and the action outcome are modeled as resulting from optimal estimates of their specific timing when relevant sensory cues and prior expectations are integrated according to causality between cues. Specifically, the shifts occur only when the “action” is inferred as causal for the subsequent effect (Legaspi and Toyoizumi 2019). In computational terms, the binding builds upon a prior belief of a causal relationship and the sensory evidence of related timing cues, independent of whether intentional action is involved. From the perspective of the Bayesian model, our increased binding of the outcome event can be viewed as reflecting an enhanced prior belief of the causal relationship between the avatar movement and the subsequent beep tone. Both our VR and control groups received identical sensory feedback in the temporal binding task, and the only difference is that the VR group had prior experience visually controlling the avatar before the post-test. The embodiment of the avatar is thus akin to an enhanced prior belief that the avatar hand is responsible for the outcome (Desantis et al. 2011; Haering and Kiesel 2012), which leads to an increased timing shift according to the Bayesian model of temporal binding (Legaspi and Toyoizumi 2019). In fact, a similar Bayesian model based on causal inference also explains the sense of bodily ownership, another component of self-consciousness, as investigated in the classical rubber hand illusion paradigm (Chancel, Ehrsson, et al. 2022). More importantly, causal beliefs about relevant ownership cues, estimated from this paradigm, are implicated in the IPS, a region often associated with cue combination, as well as the angular gyrus (Chancel, Iriye, et al. 2022). These modeling and neuroimaging work thus suggest that classical measures of self, i.e., the rubber hand illusion in the sense of bodily ownership and the temporal binding in the sense of agency, might be governed by the same causal inference mechanism with the involvement of IPL and AG.

Previous studies on VR embodiment have largely focused on how multisensory integration affects people’s self-consciousness (Slater et al. 2009; Banakou and Slater 2014). With a brief exposure to VR, people erroneously feel that they own a virtual body part or even a full virtual body (Petkova et al. 2011; Blanke et al. 2015), mislocate themselves (Ehrsson 2007), or change the perception of one’s identity (Petkova et al. 2011; Banakou et al. 2013). The common technique is to present a vivid visual representation of an avatar and match it spatiotemporally with sensory cues from other modalities, including tactile, auditory, and proprioceptive cues (Slater et al. 2009; Banakou and Slater 2014). Neuroimaging studies have shown that the premotor cortex and TPJ are key areas for bodily ownership (Bekrater-Bodmann et al. 2014; Pamplona et al. 2022) and self-location (Ionta et al. 2011; Lenggenhager et al. 2011). However, the neural correlate of SoA over a virtual body is understudied. Existing studies typically manipulated spatiotemporal mismatch between avatar and actual action (Nahab et al. 2011;

Limanowski et al. 2017) as in other embodiment studies, say, on bodily ownership. Interestingly, the neural correlates to these parametrical modulations of SoA (not necessarily about the degree of SoA) also include IPL, along with other regions like STS (Limanowski et al. 2017). Our study differed from these studies by showing that sensorimotor control experience with an avatar can lead to subsequent SoA changes over the avatar movement, whose neural correlates center at the right AG and IPL, key areas that are also associated with SoA arising from actions in real settings. Given this cluster covers high-order associative regions, we postulate that the VR embodiment effect is potentially generalizable to other tasks beyond the temporal binding. For instance, SoA arising from voluntary actions contributes to perceptual attenuation of action-induced sensory stimuli (Blakemore et al. 1998; Shergill et al. 2005) or self-other distinction (Kahl and Kopp 2018). Whether these perceptual tasks are affected by similar avatar-control experiences in VR warrants further investigation.

Our findings raised possible problems for the era of VR or metaverse. First, despite the fact that our participants did not change their self-reported SoA rating with the brief VR experience (see Supplementary Materials), it is still possible that people’s explicit judgment of SoA can be modulated by long-term VR use. Second, individuals with neurological and psychiatric disorders experience disrupted SoA and illusions in their daily lives (Frith et al. 2000; Edwards and Bhatia 2012), and even neurotypical individuals can occasionally experience faulty SoA, say, with sensory priming (Wegner and Wheatley 1999; Aarts et al. 2005). Whether certain populations’ self-consciousness is negatively affected by the experience of controlling an avatar is an important open question from the perspective of psychopathology. Third, given the observed immediate behavioral and neural effect of an embodied avatar on SoA, we expect that unintended “actions” of the avatar, accidentally caused by technical glitches in the virtual worlds, might affect the avatar owner’s sense of agency and even lead to psychological harm (Cheong 2022). These previously rare scenarios might lead to potential legal issues about how to account for the responsibility of compromising someone’s sense of agency in the metaverse.

In conclusion, the temporal binding elicited by passive observation of an embodied virtual body is subserved by the right AG and IPL, regions related to causal inference and inferential sensemaking but not directly related to motor control. In contrast, traditional motor planning areas (e.g., pre-SMA), widely observed in studies on the sense of agency arising from voluntary actions, are not implicated. These findings support the reconstructive mechanism of SoA that emphasizes retrospective processing of SoA-related cues and suggests that the experience of controlling an avatar might enhance the causal belief of avatar action and its action outcome, leading to increased temporal binding. Our behavioral and fMRI results also questioned the validity of using temporal binding as a measure of implicit SoA. Furthermore, given that people’s causal belief over an avatar’s action can be built up, both behaviorally and neurophysiologically, by a brief period of avatar control in immersive VR, how the embodiment of a virtual body affects our self-consciousness and other psychological constructs would pose as a novel problem when our populace spends increasing time in virtual or digital worlds.

## Supporting information

Supplemental results

## Acknowledgments

None

## Funding

This work was supported by the Ministry of Science and Technology of China (2021ZD0202601) and the National Natural Science Foundation of China (62061136001, 32071047, 31871102) awarded to K.W., and the National Natural Science Foundation of China (32100837) to H.Y., and the German Research Foundation (TRR 169/C8) to S.K. The funders had no role in study design, data collection and analysis, decision to publish, or preparation of the manuscript.

## Author contributions

K.W. and S.K. conceived and designed the research. Y.C., H.Y., Z.X. and K.W. implemented the methods. Y.C. executed experiments. Y.C., H.Y. and Z.X. analyzed the data. Y.C., K.W. and Y.B. wrote the original draft. X.W., H.Y., S.K., Y.B. and K.W. reviewed and edited the manuscript. K.W. supervised the study.

## Competing interests

Authors declare that they have no competing interests.

## Data and materials availability

All data needed to evaluate the conclusions in the paper are present in the paper and/or the Supplementary Materials. Data reported in this study have been made publicly available via Open Science Framework and can be accessed at https://osf.io/xnhua/.

## Notes

https://osf.io/xnhua/

### Competing Interest Statement

The authors have declared no competing interest.

### Summary of Updates

New analysis conducted; All figures revised; Author list updated.

