## Supplemental results for "Neural correlates of an illusionary sense of agency caused by virtual reality"

### Supplementary Materials

#### *Behavioral results of explicit measures*

Our previous study (Kong et al. 2017) found that the explicit SoA rating over the avatar movements did not differ between groups with and without VR exposure, nor did it correlate with the binding or the change in binding. The brief VR exposure appears insufficient to change the subjective judgments about the action attribution of the avatar hand; thus, our study focused on implicit measure, i.e., temporal binding. Nevertheless, our participants also explicitly reported their sense of embodiment for the avatar hand in the post-test (Table S1 and Fig. S1). We replicated our previous findings: we did not find a significant group difference in explicit SoA and SoO ($z$ = -.24, $p$ = .817 for explicit SoA; $z$ = -.38, $p$ = .713 for explicit SoO). Across two groups, the average scores of the control items were significantly higher than those of the agency questions ($z$ = 4.01, $p$ < .001), indicating that participants correctly followed the instructions to fill in the questionnaire. As expected, explicit SoA and SoO were positively correlated ($\rho_{43}$ = .68, $p$ < .001). However, neither of them correlated with the temporal binding effects in the post-test or the binding changes between the pre- and post-test (binding effects in the post-test: $\rho_{43}$ = .01, $p$ = .928 and $\rho_{43}$ = -.21, $p$ = .175 for explicit SoA and SoO, respectively; changes in binding effects: $\rho_{43}$ = -.01, $p$ = .925 and $\rho_{43}$ = -.05, $p$ = .722 for explicit SoA and SoO, respectively).

**Table S1. Questionnaire on embodiment evaluation for the post test**. All questions were rated on a Likert scale from 1 (strongly disagree) to 7 (strongly agree).

| Category | Questions: During the operant condition in the post-test, | Order |
|---|---|---|
| Explicit SoO | I felt like I was looking at my own hand. | 1 |
| | I felt like the avatar was part of my body. | 6 |
| | It seemed as if my hand was pressing when I saw the avatar's movement. | 5 |
| Explicit SoA | It seemed as if the avatar moved, obeying my will. | 4 |
| | It seemed as if the avatar pressed the button instead of me. | 3 |
| | If I moved my finger, I felt the virtual finger would move in the same way. | 7 |
| Control items | I felt as if the movement of the avatar had no relationship with me. | 8 |
| | It seemed as if the avatar had a will on its own. | 2 |

***Correlation analyses between fMRI and explicit measures***

For each ROI we defined, including both seven literature-based ROIs and one task-based ROI (Table 1 and Fig. 2), the correlations between the neural responses (signal in the post-test and signal change between pre-and post-test) and explicit measures (explicit SoA and SoO) were all analyzed (Table S2). Only the right CAL showed significantly positive correlations with both the explicit SoA and SoO. The right AG and IPL, showing associations with temporal bidning (Fig. 2 and Table 1), were not correlated with the explicit SoA and SoO measured in the post test. The ROI-level results, together with the lack of correlation between temporal binding and explicit ratings on the behavioral level, might suggest that that the explicit rating and temporal binding might probe agency processes at different functional and representational levels (Synofzik et al. 2008).

**Table S2. ROI-level correlations with explicit measures**.

| Location | | Correlation of explicit SoA and | | | | Correlation of explicit SoO and | | | |
|---|---|---|---|---|---|---|---|---|---|
| | | signal in post-test | | signal change | | signal in post-test | | signal change | |
| Area | H | $\rho_{43}$ | *p* | $\rho_{43}$ | *p* | $\rho_{43}$ | *p* | $\rho_{43}$ | *p* |
| **Literature-based ROIs** | | | | | | | | | |
| SMA | L | -.001 | .994 | .028 | .855 | .119 | .436 | .09 | .554 |
| Insula | L | .114 | .454 | .141 | .354 | .113 | .459 | .074 | .628 |
| CAL | R | .439 | .003 | .314 | .036 | .476 | < .001 | .362 | .015 |
| CE | R | -.033 | .829 | -.016 | .919 | .036 | .812 | -.020 | .895 |
| IPL | L | .022 | .886 | .228 | .131 | .035 | .822 | .175 | .250 |
| STG | R | .026 | .868 | .201 | .185 | .046 | .764 | .188 | .215 |
| AG | R | -.002 | .99 | .096 | .528 | -.055 | .721 | .133 | .383 |
| **Task-based ROI** | | | | | | | | | |
| IPL | R | -.016 | .915 | .227 | .134 | -.021 | .892 | .234 | .122 |

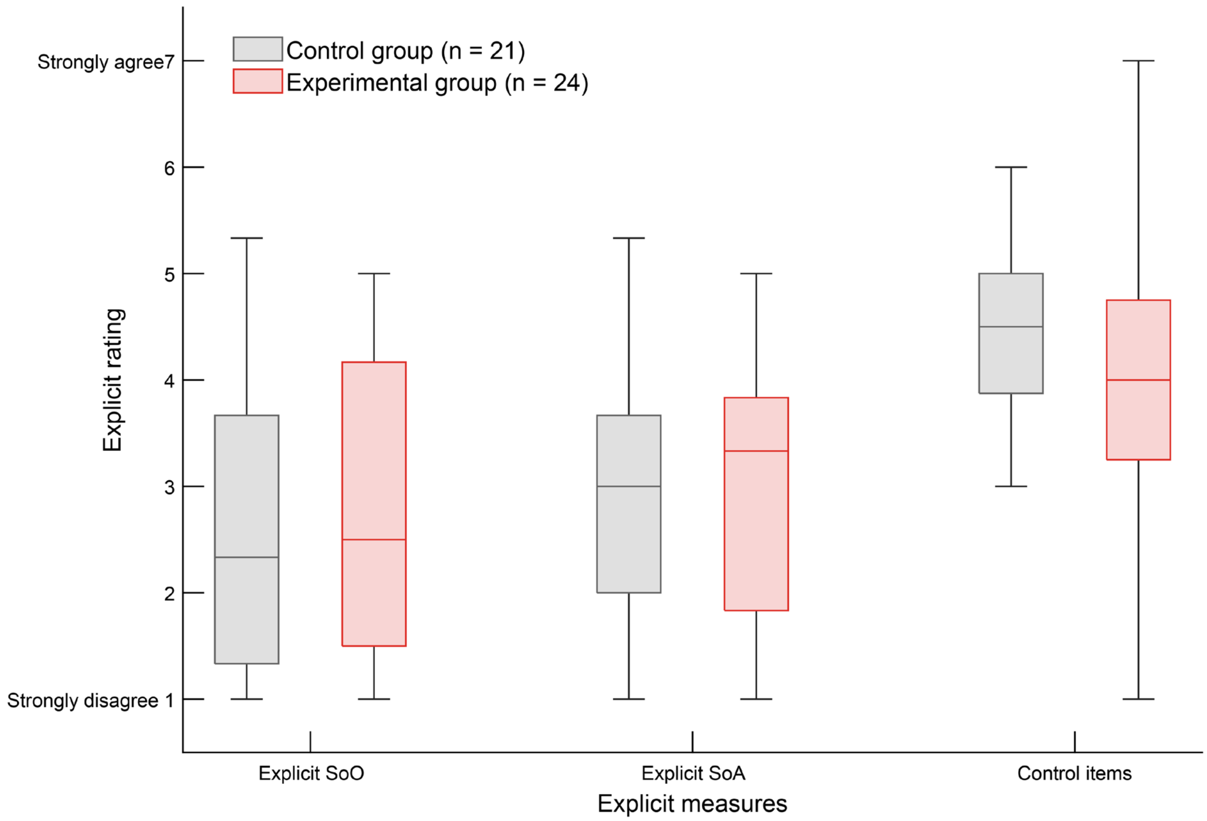


**Fig. S1. Group-based results of explicit measure of on embodiment for the post test**. Participants rated on a Likert scale from 1 (strongly disagree) to 7 (strongly agree).
